# CCG•CGG interruptions in high penetrance SCA8 families increase RAN translation and protein toxicity

**DOI:** 10.1101/2021.02.08.430311

**Authors:** Barbara A. Perez, Hannah K. Shorrock, Monica Banez-Coronel, Lauren A. Laboissonniere, Tammy Reid, Yoshio Ikeda, Kaalak Reddy, Christopher M. Gomez, Thomas Bird, Tetsuo Ashizawa, Lawrence J. Schut, Alfredo Brusco, J. Andrew Berglund, Lis F. Hasholt, Jorgen E. Nielsen, S.H. Subramony, Laura P.W. Ranum

## Abstract

Spinocerebellar ataxia type 8 (SCA8), a dominantly inherited neurodegenerative disorder caused by a CTG•CAG expansion, is unusual because most individuals that carry the mutation do not develop ataxia. To understand the variable penetrance of SCA8 we studied the molecular differences between highly penetrant families and more common sporadic cases (82%) using a large cohort of SCA8 families (N=77). We show that repeat expansion mutations from individuals with two or more affected family members have CCG•CGG interruptions at a higher frequency than sporadic SCA8 cases and that the number of CCG•CGG interruptions correlates with age at onset. At the molecular level, CCG•CGG interruptions increase RNA hairpin stability and steady state levels of SCA8 RAN polyAla and polySer proteins. Additionally, the CCG•CGG interruptions, which encode arginine interruptions in the polyGln frame increase the toxicity of the resulting proteins. In summary, CCG•CGG interruptions increase polyAla and polySer RAN protein levels, polyGln protein toxicity and disease penetrance and provide novel insight into the molecular differences between SCA8 families with high vs. low disease penetrance.

## Introduction

Spinocerebellar ataxia type 8 (SCA8) is a microsatellite expansion disorder caused by a bidirectionally transcribed CTG•CAG repeat expansion mutation within the *ATXN8OS/ATXN8* genes (Koob *et al*, 1999; Moseley *et al*, 2006). This slowly progressive cerebellar ataxia is typically characterized by ataxia, spasticity, dysarthria and nystagmus; however, extra-cerebellar features including psychiatric disturbances and developmental delays have been reported (Ayhan *et al*, 2014; Day *et al*, 2000; Juvonen *et al*, 2000; Kim *et al*, 2013; Koutsis *et al*, 2012; Lilja *et al*, 2005; Stone *et al*, 2001; Zhou *et al*, 2019). Although SCA8 is caused by a dominantly inherited mutation, patients frequently present as single affected individuals with no family history of ataxia. Despite the negative family history, asymptomatic relatives of these patients often carry the repeat expansion (Ikeda *et al*, 2004; Koob *et al.*, 1999; Moseley *et al*, 2000b; Worth *et al*, 2000). Additionally, the age of onset and clinical features of the disease vary widely among affected individuals, with onset reported from birth to 73 years of age (Day *et al.*, 2000; Felling & Barron, 2005; Ikeda *et al.*, 2004; Ikeda *et al*, 2000; Kim *et al.*, 2013; Koob *et al.*, 1999; Lilja *et al.*, 2005; Samukawa *et al*, 2019; Silveira *et al*, 2000).

Repeat associated non-AUG (RAN) proteins, which were first discovered in SCA8 and DM1 (Zu *et al*, 2011), have now been described in 11 microsatellite expansion disorders (Banez-Coronel *et al*, 2015; Banez-Coronel & Ranum, 2019; Buijsen *et al*, 2016; Goodman & Bonini, 2019; Ishiguro *et al*, 2017; McEachin *et al*, 2020; Mori *et al*, 2013; Todd *et al*, 2013; Zu *et al*, 2017; Zu *et al.*, 2011). These repetitive proteins expressed by repeat associated non-ATG (RAN) translation accumulate in affected brain regions in SCA8 patients (Ayhan *et al*, 2018; Zu *et al.*, 2011). RAN translation is a process in which transcripts containing repeat expansions express proteins in multiple reading frames without the requirement of AUG- or AUG-like close-cognate initiation codons (Banez-Coronel & Ranum, 2019; Cleary *et al*, 2018; Nguyen *et al*, 2019; Zu *et al.*, 2011). The presence of RAN and ATG-initiated expansion proteins has been previously reported in human SCA8 autopsy brains and SCA8 BAC transgenic mice (Ayhan *et al.*, 2018; Moseley *et al.*, 2006; Zu *et al.*, 2011). Both ATG-initiated poly-glutamine (polyGln) and RAN poly-Alanine (polyAla) have been found in Purkinje cells (Moseley *et al.*, 2006; Zu *et al.*, 2011) and polyGln and RAN poly-Serine (polySer) proteins in the hippocampus, pons and frontal cortex (Ayhan *et al.*, 2018). Additionally, polySer aggregates are found in the cerebellar white matter and brainstem nuclei where they are associated with demyelination, axonal degeneration, increased astrogliosis and a reduction in the number of mature oligodendrocytes (Ayhan *et al.*, 2018).

In contrast to other SCAs, SCA8 is unusual in that there is markedly reduced penetrance. The reduced penetrance is consistent with the detection of SCA8 expansions in the general population and the variable age of onset (Ikeda *et al.*, 2004; Koob *et al.*, 1999; Stevanin *et al*, 2000; Worth *et al.*, 2000), suggesting that genetic and/or environmental modifiers affect the onset and penetrance of SCA8. One potential genetic modifier that may affect disease penetrance in SCA8 is the presence of repeat interruptions. The presence of CCG, CTA, CTC, CCA and CTT interruptions in the CTG repeat expansion in SCA8 has previously been reported (Hu *et al*, 2017; Moseley *et al.*, 2000b). These interruptions can vary in number, configuration and the position within the repeat tract. Interestingly, one to four CCG interruptions were detected in multiple configurations among affected members of a large highly penetrant SCA8 family (MN-A) and the number of interruptions often increases when passed from one generation to the next (Moseley *et al.*, 2000b).

Repeat interruptions have been reported to have different modifying effects in a number of other microsatellite disorders. For several of these disorders (SCA1, SCA2 and FXS), sequence interruptions appear to stabilize repeat tracts found on unexpanded alleles, and the loss of interruptions predisposes repeat tracts to expand above the pathogenic threshold (Chung *et al*, 1993; Gunter *et al*, 1998; Imbert *et al*, 1996; Kunst & Warren, 1994; Pulst *et al*, 1996; Sanpei *et al*, 1996). In other cases, interruptions do not stabilize normal alleles but are found on expanded alleles and are associated with changes in disease presentation. For example, CAA interruptions on expanded alleles are associated with later ages of onset in SCA2 (Sobczak & Krzyzosiak, 2005) and Huntington disease (Genetic Modifiers of Huntington’s Disease, 2019; Wright *et al*, 2019). In SCA10, patients with ATCCT interruptions are prone to seizures (McFarland *et al*, 2014) and in DM1 CCG and GGC interruptions are found in patients with peripheral neuropathy (Braida *et al*, 2010), but the molecular basis for these effects is unclear.

Here, we show that CCG•CGG interruptions are preferentially found on SCA8 alleles in families with increased disease penetrance and that age of onset is inversely correlated with the number of interruptions and not repeat length. Molecular studies show CCG•CGG interruptions increase polyAla and polySer RAN protein levels and the toxicity of the resulting arginine interrupted polyGln expansion proteins. Our demonstration that CCG•CGG interruptions increase RAN protein levels and polyGln protein toxicity and are found in families with increased disease penetrance provides novel molecular insight into the variable penetrance and risk of developing SCA8.

## Results

### Most SCA8 patients have no family history of ataxia

To investigate the effects of sequence interruptions in SCA8, we performed a detailed genetic evaluation of expanded SCA8 alleles from a large cohort of SCA8 families (N=77) including 199 expansion carriers (n=111 affected, n=88 asymptomatic). Disease onset ranged from birth to 79 years with an average age of onset of 33.7 years (Table EV1). Although the mutation is transmitted in an autosomal dominant pattern, surprisingly 82% (63/77) of these families had sporadic ataxia with no family history of disease, 5% (4/77) had family histories that appeared recessive and only 13% (10/77) showed the expected autosomal dominant inheritance pattern (Fig 1A). Interestingly, four of the sporadic and two familial cases are homozygous and have two expanded alleles. These data and previous reports of expansion alleles in unaffected family members and in the general population (Cellini *et al*, 2001; Ikeda *et al.*, 2004; Moseley *et al*, 2000a; Stevanin *et al.*, 2000; Worth *et al.*, 2000; Zeman *et al*, 2004) highlight the need to understand the molecular basis of the variable penetrance found in SCA8 families.

**Figure 1.**
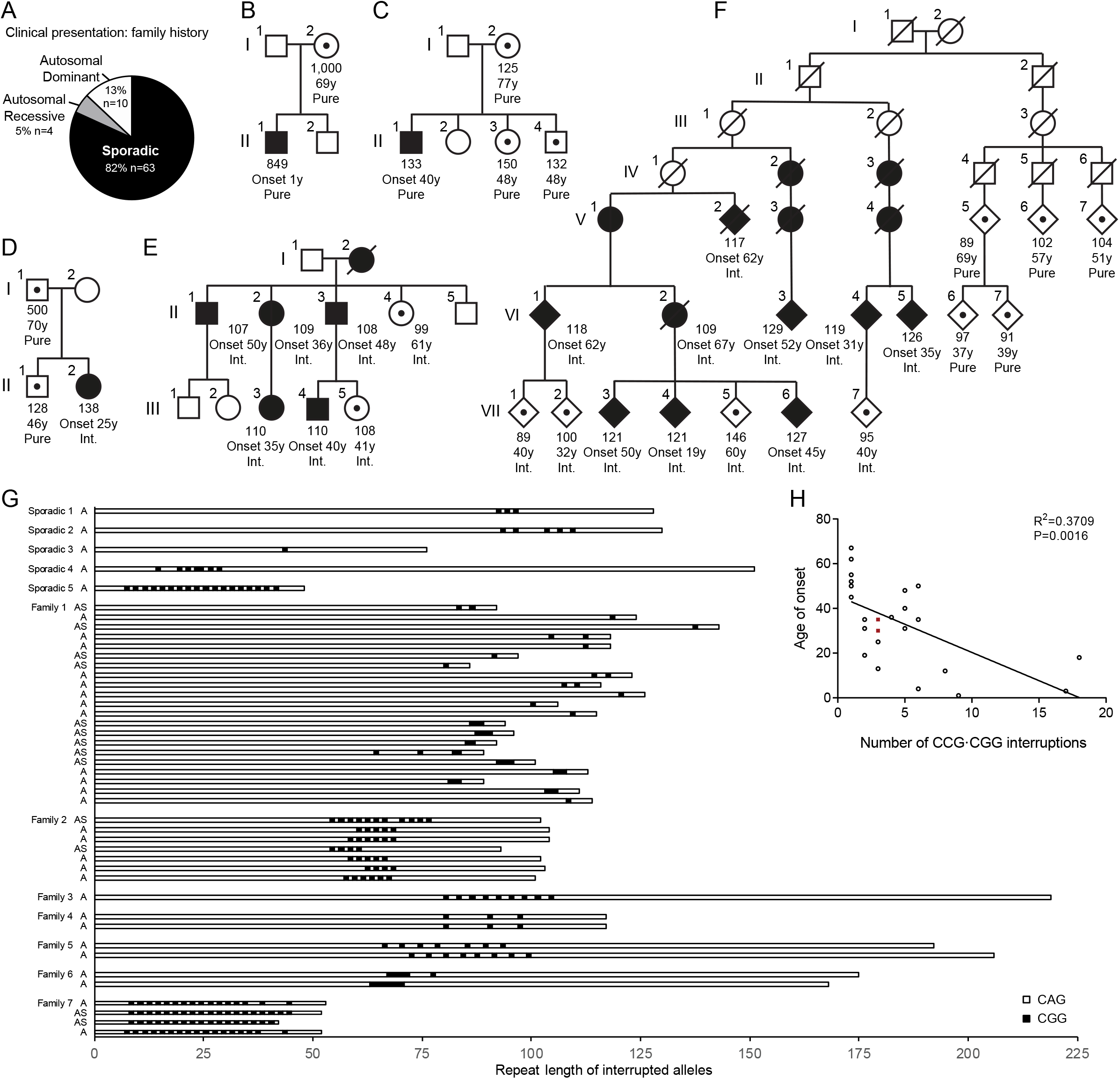
SCA8 alleles with CCG•CGG interruptions are found in families with high disease penetrance. A Summary of family history of SCA8 disease presentation in the clinic for the 77 SCA8 families in our cohort. B-F Pedigrees of SCA8 families: squares represent males, circles represent females, diamonds mask gender. Filled symbols represent affected individuals, symbols with inner black dot represent asymptomatic expansion carriers, open symbols represent individuals with non-expanded alleles, diagonal line indicates a deceased individual. Combined repeat number of expanded alleles, age (y-years old) at onset (Onset) or age still asymptomatic, and interruption status (Pure or Int. [CCG•CGG interrupted]) are noted below the symbols. (F) Abbreviated pedigree, for complete pedigree see Koob *et al.*, 1999. G SCA8 allele configurations in the CAG direction as determined by sequencing. Family or individual and affected status indicated on left: Sporadic 1 - figure 1D, Family 1 - figure 1F, Family 2 - figure 1E; A – affected, AS – Asymptomatic; CGG interruptions are represented by black boxes. H Age of onset correlates with the number of CCG•CGG interruptions, n=24, p=0.0016. Red squares indicate the average expansion size for individuals with two expanded alleles, individual allele repeat lengths are: 84/114, 92/100. G, H Individuals identified as having CCG•CGG interruptions by restriction digest are not included.

### SCA8 repeat length does not correlate with age of onset or predict disease status

Similar to previous reports (Ayhan *et al.*, 2014; Ikeda *et al.*, 2004; Juvonen *et al.*, 2000; Zeman *et al.*, 2004), we found no correlation in the number of SCA8 repeats and age of onset (Fig EV1A), no significant difference in repeat length between affected patients (median: 113 repeats) and asymptomatic carriers (median: 98 repeats; p=0.0672; Table EV1) and a wide and overlapping range of repeat lengths in affected (54-1455) and asymptomatic individuals (52-1000) (Fig EV1B; Table EV1). The lack of correlation of repeat length and disease status is often seen in individual SCA8 families. For example, in Fig 1B, individual I-2 carries an expansion of 1000 repeats yet remains asymptomatic, while individual II-1 has an expansion of 849 repeats and presented with disease at one year of age. Similarly, in Fig 1C, individual II-1 presented with disease at age 40 with 133 combined repeats while her mother and two siblings, who carry SCA8 expansions of similar lengths, remain asymptomatic. Taken together, these data provide additional evidence that repeat length is not a reliable predictor of disease or age of onset and suggest other genetic or environmental modifiers contribute to the variable penetrance of SCA8.

A potential genetic modifier of SCA8 is the presence of interruptions within the CAG repeat expansion. In Fig 1D a 25-year-old female (II-2), with no family history of SCA8, has an expansion mutation containing three *de novo* CGG interruptions [(CAG)_91_(CAGCGG)_3_(CAG)_31_(TAG)_10_]. These interruptions were not found in her asymptomatic 70-year-old father (I-1; confirmed pure by MspA1I digest) or 46-year-old brother (II-1; (CAG)_118_(TAG)_10_; Fig 1D). The observation that the only affected individual in this family has CCG•CGG interruptions combined with the previously reported CCG•CGG interruptions in affected members of an unusually large SCA8 kindred (Moseley *et al.*, 2000b), suggests that CCG•CGG interruptions are associated with increased disease penetrance.

### CCG•CGG interruptions increase disease penetrance and inversely correlate with age of onset

To better understand the effects of CCG•CGG interruptions on disease penetrance, we compared the sequences of SCA8 expansion alleles in families with high (≥3 affected) versus low disease penetrance. The seven-generation MN-A family (Day *et al.*, 2000; Koob *et al.*, 1999), the largest SCA8 family reported to date, has a much higher disease penetrance than most SCA8 families (Ikeda *et al.*, 2004) and CCG•CGG interruptions were reported in all affected individuals (Moseley *et al.*, 2000b). Additional analyses of this family show CCG•CGG interruptions are found in the high but not a newly identified low penetrance branch of this family. The left family branch shows an autosomal dominant inheritance pattern (onset 19-74 years; Fig 1F) while members of the extended right branch have pure CTG•CAG expansions and no affected individuals. In a second newly identified multigenerational family, all six affected individuals (onset 35-50 years; Fig 1E) have CCG•CGG interruptions. These interruptions were also identified in individual II-4 who was not affected at the time of examination but subsequently showed signs of ataxia and in individual III-5 who was asymptomatic at age 41 (Fig 1E). CCG•CGG interruptions were found at a higher frequency in families with multiple affected individuals: 100% (5/5) of families with three or more affected individuals, 28.6% (2/7) of families with two affected individuals and 13.9% (5/36) of sporadic cases. Overall, CCG•CGG interruptions were found at a higher frequency in SCA8 families with 2 or more affected members compared to sporadic cases (n=48; p=0.0047; Table 1) and among affected individuals compared to asymptomatic carriers (n=132; p=0.0299; Table 1. While the position, configuration and number of CCG•CGG interruptions varies widely among SCA8 families (Fig 1G), the number of CCG•CGG interruptions is inversely correlated with, and accounts for 37% of the variation in age of onset (R^2^=0.3709; p=0.0016; Fig 1H).

**Table 1.** CGG interruptions are associated with increased disease penetrance in SCA8. P-value was calculated using Fisher’s exact test to assess the relationship between disease penetrance and CGG interruptions (n=48 families; n=132 expansion carriers). An additional n=10 families, representing n=19 expansion carriers, were sequenced and found to carry different interruptions.

**SCA8**
|  | Pure | CGG interrupted | p-value |
| --- | --- | --- | --- |
| Apparent sporadic patients | 31 | 5 | 0.0047 |
| Families with 2+ affected individuals | 5 | 7 |  |
| Affected individuals | 41 | 33 | 0.0299 |
| Asymptomatic individuals | 43 | 15 |  |

Taken together, these data demonstrate that CCG•CGG interruptions increase disease penetrance and that the number of interruptions, and not repeat length, is inversely correlated with age at onset in SCA8.

### CCG•CGG interruptions increase the toxicity of SCA8 CAG•CTG repeat expansions

To better understand the molecular effects of interrupted alleles we examined if constructs containing CCG•CGG interruptions are more toxic to cells than pure expansion constructs. T98 glial cells were transfected with length-matched constructs containing pure or interrupted expansions cloned from patient DNA and expressed in the CAG direction (Fig 2A). Interrupted expansions were cloned from individuals from the high-penetrance multigeneration families shown in Fig 1F (Int.95) and Fig 1E (Int.102). Int.95 contains an overall CAG repeat length of 95 with 4 consecutive CGG interruptions near the 3’ end, followed by 3 TAGs which were found in this patient [(CAG)_86_(CGG)_4_(CAG)_5_(TAG)_3_]. Int.102 contains 4 mixed CAGCGG interruptions in the middle of the CAG repeat for a total of 102 interrupted CAGs followed by 6 TAGs [(CAG)_63_(CGGCAG)_4_(CAG)_31_(TAG)_6_] (Fig 2A). Cells expressing these interrupted constructs showed increased death (26.9%, p<0.05 – Int.95 vs Pure 96; 23.5%, p<0.05 – Int.102 vs Pure 104; Fig 2B) and decreased viability (16.5%, p<0.05 – Int.95 vs Pure 96; 15.6%, p<0.05 – Int.102 vs Pure 104; Fig. 2C) compared to length matched uninterrupted repeats. These effects cannot be explained by differences in RNA levels which did not differ in Pure-96 and Int-95 transfected cells and were actually lower in Int-102 vs Pure 104 transfected cells (Fig EV2). Taken together, these data indicate that CGG interruptions increase the toxicity of CAG repeats independent of RNA levels.

**Figure 2.**
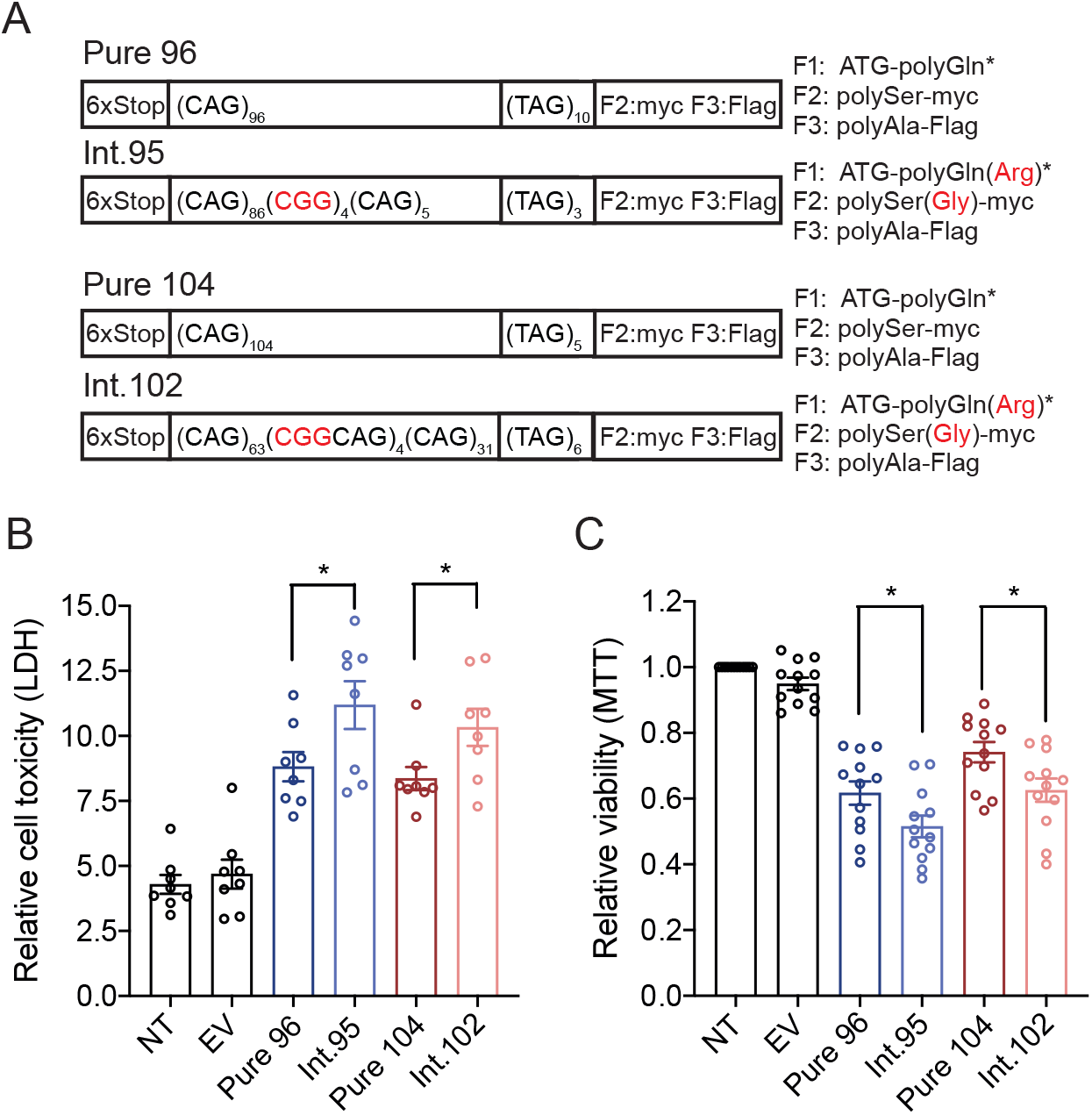
Clustered and interspersed CCG•CGG interruptions increase toxicity of CTG•CAG expansions. A Schematic diagram of constructs used to express patient-derived pure and interrupted SCA8 repeat tracts with predicted protein products and C-terminal epitope tags. * Due to TAG encoded stop codons polyGln proteins do not contain epitope tags. CGG interruptions and the encoded interruption amino acids are indicated in red. B, C Cell death measured by lactase dehydrogenase (LDH) assay (B) and cell viability measured by 3-(4,5-dimethyl-thiazol-. 2-yl)-2,5-diphenyl tetrazolium bromide (MTT) assay (C) in T98 cells 42 hrs post-transfection of pure and interrupted SCA8 repeat tracts; LDH n=8, MTT n=12, * p < 0.05, NT not transfected, EV - empty vector (pcDNA3.1-6S-3T).

### Arginine-encoding CGG interruptions increase toxicity of polyGln proteins

Next, we tested the hypothesis that CGG interruptions increase the toxicity of expanded alleles by affecting RAN and polyglutamine proteins expressed from the CAG repeat. First, we examined if the arginine interruptions in the polyGln(Arg) proteins increase their toxicity compared to pure polyGln proteins. To perform these experiments, we generated minigene constructs to express polyGln and polyGln(Arg) using non-hairpin forming alternative codons (Fig 3A). This enables the toxicity of pure and interrupted proteins to be assessed individually and independent of possible effects from CAG expansion RNAs and RAN proteins. We focused these experiments on pure and interrupted polyGln proteins because non-hairpin forming alternative codons are available for both Gln and Arg. Transient transfections in T98 cells show that interrupted polyGln(Arg) proteins expressed with alternative codons increased cell death by 25% (p<0.05; Fig 3B) and decreased cell viability by 10% compared to pure polyGln proteins (p<0.05; Fig 3C), independent of RNA levels (Fig EV3A). Protein blot and immunofluorescence analyses show that the pure and arginine interrupted polyGln proteins have different properties. For example, the interrupted polyGln(Arg) proteins migrate further into the gel (Fig 3D, EV3B) and show droplet-like nuclear staining not found with pure polyGln proteins (Fig 3G). These changes may contribute to the increased toxicity of the polyGln(Arg) proteins. Surprisingly, substantially less polyGln(Arg) compared to pure polyGln protein was detected by 1C2 antibody (Fig 3D-F). This may be caused by reduced affinity of the 1C2 antibody for the interrupted protein or incomplete extraction of polyGln(Arg) proteins from nuclear aggregates. Taken together, these data demonstrate that arginine interruptions increase the toxicity of polyGln expansion proteins and that the increased toxicity of the interrupted polyGln(Arg) proteins is independent of possible CAG RNA gain-of-function or RAN protein.

**Figure 3.**
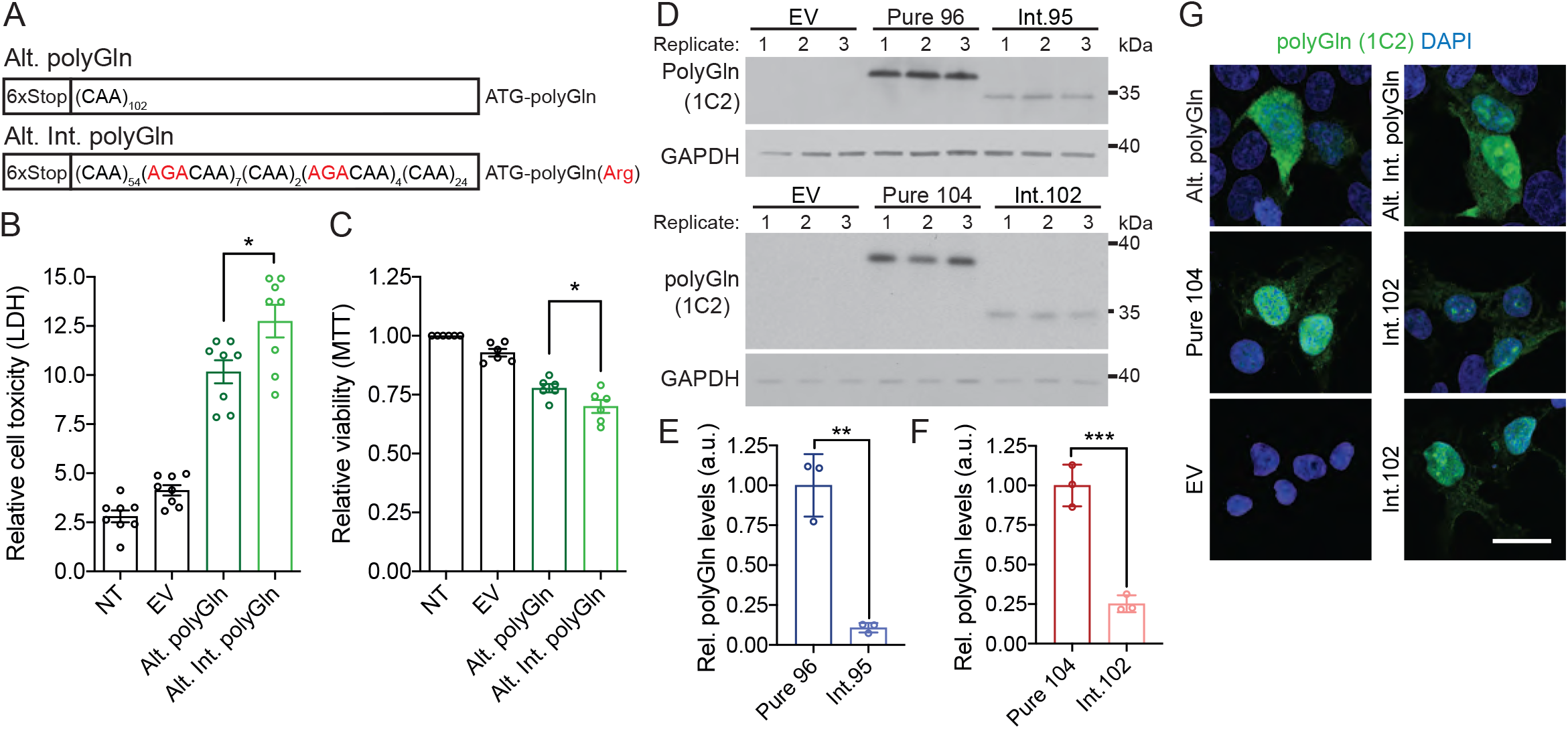
Arginine-encoding CGG interruptions increase toxicity of ATXN8 polyGln proteins. A Schematic diagram of alternative-codon constructs expressing pure and interrupted polyGln proteins. B, C Cell death measured by LDH assay (B) and cell viability measured by MTT assay (C) in T98 cells 42 hrs posttransfection of alternative-codon polyGln constructs, LDH n=8, MTT n=6, data are presented as mean ± SEM. NT: non-transfected; EV – empty vector (pcDNA3.1-6S-3T); * p < 0.05. D-F Western blot (D) and densitometry quantification (E, F) of polyGln proteins in HEK293T cells detected by 1C2 antibody from interrupted and pure CAG repeat tracts; EV: empty vector (pcDNA3.1-6S-3T), GAPDH as loading control; n=3, ** p<0.01, *** p<0.001, data presented as mean ± SD. G Immunofluorescence of polyGln expressed from Alt. polyGln, Alt. Int. polyGln, Pure 104 and Int.102 constructs in HEK293T cells, scale bar: 20μ m; EV-empty vector (pcDNA3.1-6S-3T).

### CGG interruptions increase polyAla and polySer RAN protein levels

Next, we examined the effects of the CGG interruptions on polySer and polyAla RAN proteins. Transient transfections with interrupted and pure repeat constructs show CGG interruptions substantially increase steady state levels of polySer and polyAla RAN proteins (Fig 4). In the polySer reading frame, the GGC interruptions produce a polySer protein with glycine interruptions, polySer(Gly). Dot blot analyses showed 93.8% higher levels of interrupted RAN polySer(Gly) compared to pure RAN polySer proteins (p<0.01; Fig 4A, B) and immunofluorescence showed RAN polySer(Gly) proteins form globular or clustered aggregates compared to punctate aggregates formed by pure polySer RAN proteins (Fig 4C).

**Figure 4.**
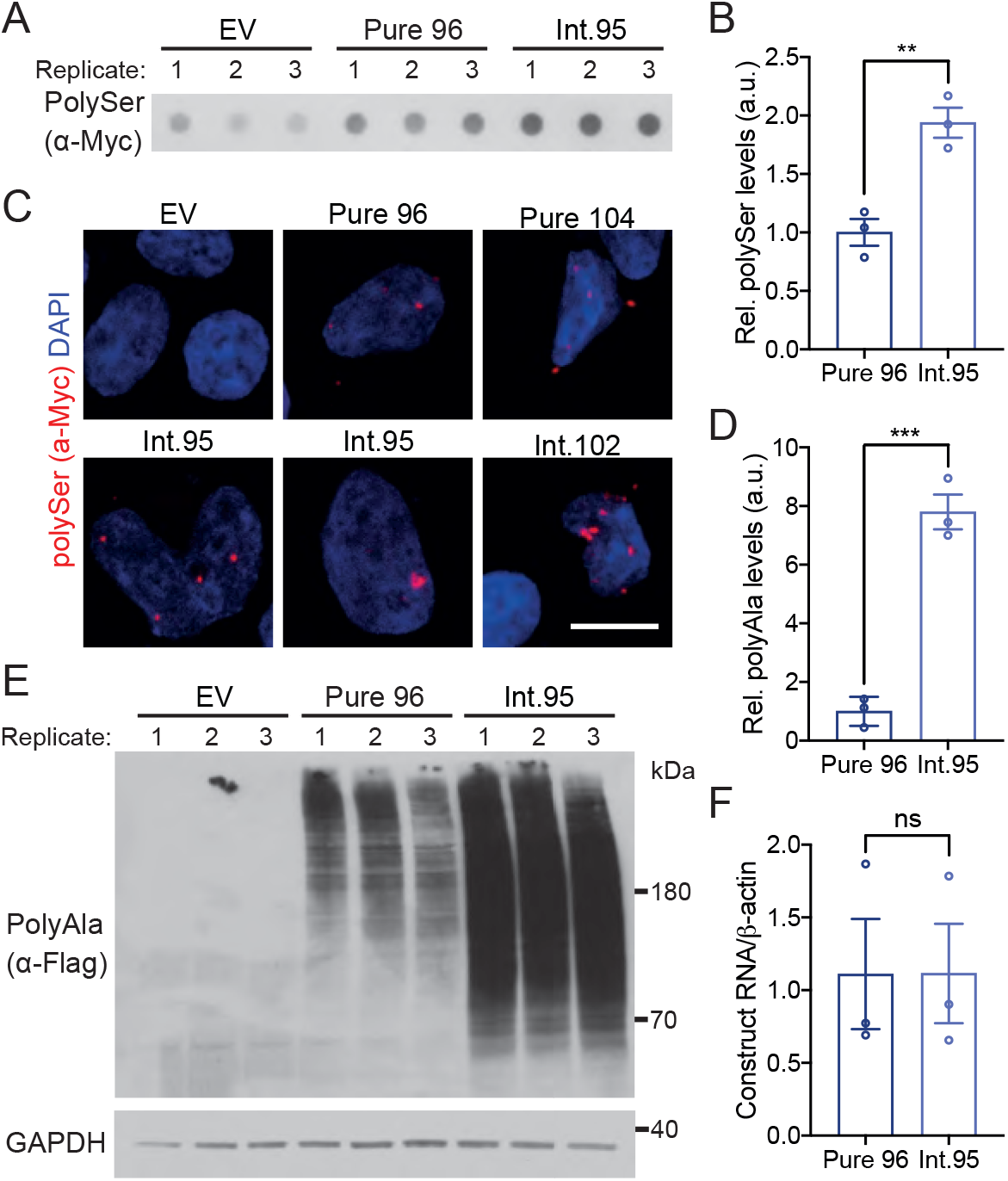
CGG interruptions increase RAN polySer and RAN polyAla protein steady state levels. A-B Protein blotting (A) and densitometry quantification (B) of polySer RAN proteins in HEK293T cells from interrupted (construct Int.95) and pure (construct Pure 96) CAG repeat tracts. EV: empty vector (pcDNA3.1-6S-3T), n=3, ** p<0.01, data are presented as mean ± SEM. C Immunofluorescence of RAN polySer protein aggregates from CGG interrupted and pure CAG repeat tracts in HEK293T cells; scale bar: 10μ m. D-E Protein blotting (E) and densitometry quantification (D) of polyAla RAN proteins; n=3, *** p<0.001, data are presented as mean ± SD. F qRT-PCR of Pure 96 and Int.95 construct transcript levels; n=3; p=0.9942, data are presented as mean ± SEM.

Protein blots showed even higher increases (7.8-fold) in steady state levels of polyAla RAN proteins expressed from interrupted (Int. 95) compared to pure (Pure 96) CAG repeats (p<0.001; Fig 4D, E). Transfections with constructs containing interspersed CGG interruptions (Int. 102) showed similar polyAla increases (2.8 fold) compared to size comparable pure repeats (Pure 104) (p<0.01; Fig EV4A, B). The increases in polyAla protein levels did not show overt changes in cellular localization of the polyAla proteins (Fig EV4C) and were not caused by changes in RNA levels (Fig 4F, EV4D).

Taken together, these data show CGG interruptions increase steady state levels of polySer and polyAla RAN proteins independent of RNA levels. Additionally, the fact that pure polyAla proteins are expressed from both interrupted and pure CAG expansions indicates that the increase in steady state levels of polyAla RAN proteins is not caused by changes in the nature or stability of the polyAla protein.

### CGG interruptions increase stability of CAG expansion transcript secondary structure

RAN translation is favored by repeat length and RNA structure (Banez-Coronel *et al.*, 2015; Wang *et al*, 2019; Zu *et al.*, 2011; Zu *et al*, 2013) and RNA hairpin stability is known to increase with repeat length (Napierala *et al*, 2005; Wang *et al.*, 2019). CGG interruptions increase the steady state levels of polyAla without altering the amino acid sequence, suggesting that the increased levels of RAN proteins expressed from interrupted alleles are caused by changes in RNA structure or stability. Consistent with this hypothesis, UV melting analyses of RNA oligos with CGG interruptions required higher melting temperatures than oligos with pure repeats (Fig 5A, EV5). Additionally, computational predictions using *m*-fold (Zuker, 2003) of short RNAs show increased stability with the presence of CGG interruptions (Fig 5B, EV6A). Next, we examined the stability of interrupted alleles found in patients. We used *m*-fold to compare the stability of several highly interrupted full-length CAG-repeat tracts from patients (48-53 repeats), with length-matched pure repeats. Results from these analyses show that the multiple predicted hairpin structures, including branched structures, are more stable for alleles containing CGG interrupted CAG repeats compared to length-matched pure CAGs (Fig 5C, EV6B). Both interruption number and configuration influence RNA structural stability in computational (Fig 5B) and UV-melting (Fig 5A) analyses. Taken together, these data are consistent with a model in which increased stability of CGG-interrupted expansion transcript secondary structures increases RAN translation.

**Figure 5.**
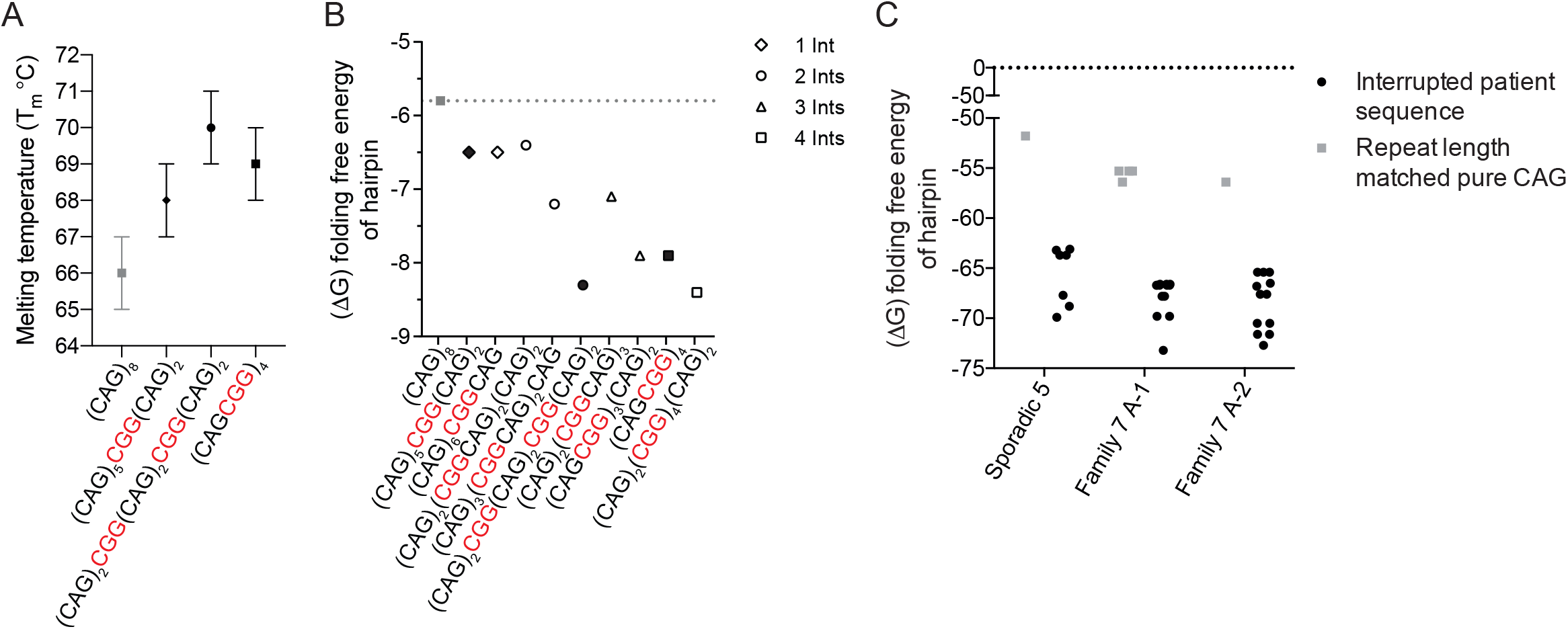
CGG interruptions increase stability of CAG repeat RNA hairpins. A Absorbance of each RNA substrate at 260nm monitored between 25°C and 95°C, recorded at 1°C intervals; n=3 UV melting curves per RNA substrate at a concentration of 2μ M in 1xDPBS without calcium or magnesium. B The folding free energy (ΔG) of hairpin structures for pure CAG and CGG interrupted repeat tracts for different interruption configurations, as predicted by m-fold (Zuker, 2003). Filled symbols represent sequences used for UV melting analyses. C The folding free energy (ΔG) of hairpin structures for SCA8 patient repeat expansions (Figure 1G) and pure repeat tracts of the same length, as predicted by m-fold. Patient alleles are as follows: 48 repeats in length -(CAG)_7_(CGGCAG)_18_(CAG)_5_; 53 repeats in length - (CAG)_8_(CGGCAG)_14_(CAG)_2_CGG(CAG)_5_CGG(CAG)_8_; and 52 repeats in length – (CAG)_7_(CGGCAG)_16_(CAG)_4_CGG(CAG)_8_. Each symbol represents a single predicted hairpin structure; multiple hairpin structures, including branched hairpins, are predicted for SCA8 patient alleles and (CAG)_53_ (Zuker, 2003).

## Discussion

The markedly reduced penetrance is one of the most puzzling features of SCA8 (Ikeda *et al.*, 2004; Koob *et al.*, 1999; Stevanin *et al.*, 2000; Worth *et al.*, 2000). Here we show that 82% of SCA8 families in a large cohort of SCA8 families have only a single affected individual, even though the repeat expansion mutation is inherited in an autosomal dominant manner. A much smaller percentage of families (13%) showed the expected autosomal dominant pattern of disease. Here we show CCG•CGG interruptions in the CTG•CAG repeat tract are found at a higher frequency in families with multiple affected individuals and that the number of CCG•CGG interruptions, and not repeat length, correlates with age at onset. Cell culture studies show CAG expansions with CGG interruptions are more toxic than pure repeats. At the protein level, CGG interruptions within the CAG repeat tract increase steady state levels of the SCA8 RAN polyAla and polySer proteins. This observation is consistent with the increased stability of RNA structures predicted on CGG interrupted alleles. It will be interesting in future work to understand if PKR activation, which is activated by structured microsatellite RNAs (Edery *et al*, 1989; Tian *et al*, 2000; Zu *et al*, 2020) and which has been recently shown to be a major driver of RAN translation (Zu *et al.*, 2020), is also increased by CGG interruptions. Additionally, CGG interruptions introduce arginine amino acids into the polyGln proteins which increases their toxicity. Taken together, these data demonstrate that CCG•CGG interruptions act as *cis*-modifiers of SCA8 and provide a molecular explanation for the dramatic variations in disease penetrance among SCA8 families.

We found CCG•CGG interruptions on expanded alleles in all families in our cohort with three or more cases of SCA8. CCG•CGG interruptions were also identified in sporadic SCA8 cases, but at a lower frequency. Additionally, we confirm that repeat length in SCA8 is a poor predictor of disease penetrance (Ikeda *et al.*, 2004; Stevanin *et al.*, 2000; Worth *et al.*, 2000). Taken together, these data indicate that the inclusion of sequence information during genetic testing, specifically the presence or absence of CCG•CGG interruptions, will provide patients and families with additional information relevant to disease penetrance. Sequence analyses will also further our understanding of the role of additional types of interruptions on disease penetrance in SCA8 and help identify the causes of high penetrance in other large SCA8 families in the literature for which the expansion sequences are unknown (Cintra *et al*, 2017). Additionally, we identify SCA8 patients with shorter repeat expansions than have been previously reported, expanding the range of repeats found in individuals affected with ataxia to 54-1455 repeats.

Here we show that the polyGln proteins produced from interrupted SCA8 transcripts are more toxic and that steady state levels of the RAN polyAla and polySer proteins are increased. Together it is possible that in SCA8 patients, the CGG interrupted repeat expansions increase overall cellular toxicity and RAN protein load which may in turn exacerbate the associated pathologies, including white matter defects (Ayhan *et al.*, 2018), in SCA8. However, while the data presented here provide insight into possible molecular consequences of the CCG•CGG interruptions in SCA8 repeat expansions, further detailed analyses in patient cell lines and postmortem tissue, which are currently very limited for CCG•CGG interrupted expansions, will be necessary to fully understand the pathological consequences of the CCG•CGG interruptions. In addition, directly comparing tissues and cell lines from SCA8 patients with pure and CCG•CGG interrupted repeat expansions will help to inform our understanding of the contribution of repeat expansion proteins to disease.

There is a growing body of evidence that structured RNAs, including RNA hairpins favor efficient RAN translation (Banez-Coronel *et al.*, 2015; Wang *et al.*, 2019; Zu *et al.*, 2011). RAN translation has also been shown to be more efficient with longer repeat lengths and longer repeats which increase the structural stability of RNA secondary structures (Banez-Coronel *et al.*, 2015; Napierala *et al.*, 2005; Wang *et al.*, 2019; Zu *et al.*, 2011; Zu *et al.*, 2013). Our data extend these results and show that CGG interruptions, which increase the stability of RNA hairpins, also lead to elevated levels of RAN proteins and independently show that increasing RNA stability without altering repeat tract length increases RAN translation. Additionally, the increased stability of RNA secondary structures containing CGG interruptions could also lead to increased toxicity through RNA gain-of-function mechanisms (Daughters *et al*, 2009) possibly by the changes in the sequestration of known and novel RNA binding proteins by SCA8 expansion transcripts.

While additional types of AT-rich sequence interruptions (e.g. CTT•AAG, CCA•TGG, CTA•TAG) have been reported in SCA8 (Hu *et al.*, 2017; Moseley *et al.*, 2000b), the lack of highly penetrant SCA8 families with AT-rich interruptions (Moseley *et al.*, 2000b) makes it unlikely that they increase disease penetrance in a manner similar to CGG repeats. This is consistent with the prediction that AT-rich interruptions decrease RNA structural stability of CAG expansion transcripts in contrast to CGGs, which increase RNA stability. A small number of sporadic cases are homozygous for the expansions suggesting the presence of two SCA8 expansion alleles may also increase disease penetrance (Fig EV1A). The fact that SCA8 is also found with reduced penetrance in patients with single uninterrupted expansion mutations suggest that, similar to other neurodegenerative diseases, *trans*-genetic modifiers and environmental factors are also likely to contribute to disease (Hosseinibarkooie *et al*, 2017; Mo *et al*, 2015).

In summary, CCG•CGG interruptions within the SCA8 CAG repeat tract are associated with increased penetrance in SCA8 families. At the molecular level CCG•CGG interruptions increase RNA stability and levels of polyAla and polySer RAN proteins. Additionally, CCG•CGG interruptions encode alternative amino acids that increase the toxicity and change the molecular properties of the resulting polyGln(Arg) proteins. Taken together, these data provide novel insight into the molecular mechanisms affecting disease penetrance in SCA8.

## Materials and Methods

### Research participants

Informed consent was acquired from all participants in accordance to the Human Subjects Committee at the University of Minnesota, the Institutional Review Board (IRB) at the University of Florida, or the equivalent office at collaborators’ institutions. A board-certified neurologist identified SCA8 probands on clinical examination and interested patients were enrolled into the research study. Family history of ataxia was assessed by questionnaire and patients were encouraged to inform affected and unaffected relatives of the research study; volunteers were enrolled into the study. Samples were collected from 77 independent families.

### Genetic analysis of SCA8 repeat expansions

Genomic DNA (gDNA) was extracted from peripheral blood lymphocytes using FlexiGene DNA kit (QIAGEN). The number of combined CTG•CAG repeats at the SCA8 locus was determined by PCR across the repeat using CAG-1F (5’ TTT GAG AAA GGC TTG TGA GGA 3’) and CAG-1R (5’ TCT GTT GGC TGA AGC CCT AT 3’) primers. PCR bands were extracted using Wizard SV Gel and PCR Clean-Up System (Promega) and, when possible, sent for direct DNA sequencing using nested primers CAG-3F (5’ GGC TTG TGA GGA CTG AGA ATG 3’) and CAG-3R (5’ GAA GCC CTA TTC CCA ATT CC 3’). Expansions too large for direct sequence (approximately >250 repeats) were digested with MspA1I (New England Biolabs) which ambiguously digests the PCR products containing either CGG or CTG interruptions in the CAG direction of the repeat tract. This method does not provide the sequence configuration. If the expansion size was too large to perform PCR across the repeat or we were unable to draw blood from the subject, the repeat length was estimated by a commercial diagnostic company. Families found to have non-CGG interruptions were excluded from analysis of CGG interruptions and disease penetrance (n=58 families sequenced in total).

### cDNA constructs

To generate patient derived pure and interrupted SCA8 expansion constructs for molecular characterization of CGG interruptions, a region containing the *ATXN8* open reading frame was PCR amplified from patients’ gDNA using primers SCA8-F3-Kpn1 (5’ TTG GTA CCT TTG AGA AAG GCT TGT GAG GAC TGA GAA TG 3’) and SCA8-R4-EcoRI (5’ GCG AAT TCG GTC CTT CAT GTT AGA AAA CCT GGC T 3’). The PCR fragment was cloned in the CAG direction into the pcDNA3.1-6S-3T vector which has six stop cassette (two stop codons per reading frame) upstream of the repeat and a unique C-terminal tag in each reading frame (Zu *et al.*, 2011). Due to the TAG repeat tract encoding for multiple stop codons after the CAG repeat stretch, there is no C-terminal tag in the CAG frame. Additionally, construct names denote the total CAG tract length which, due to repeat instability during cloning, may not be the same total tract length as the patient alleles used to clone the repeat sequences.

To assess toxicity of polyGln proteins, ATG-initiated non-hairpin forming alternative codon minigenes were synthesized by IDT Technologies and subcloned into the pcDNA3.1-6S-3T vector. PolyGln is encoded by CAA repeats with AGA-encoded Arginine interruptions to generate the Alt. polyGln and Alt. Int. polyGln constructs (Fig 6A). It is not possible to model polyAla or polySer proteins using this system as no non-hairpin forming alternative codons exist for alanine or glycine.

### Cell culture and transfections

HEK293T or T98 cells were cultured in DMEM medium (Corning) supplemented with 10% fetal bovine serum (FBS) (Gibco) and 1X Penicillin-Streptomycin (Gibco). Plasmid transfections were performed using Lipofectamine 2000 (Invitrogen), according to the manufacturer’s instructions. Plasmid transfection amounts were optimized for each set of constructs used for toxicity assays.

### Toxicity and viability assays

Cell toxicity and viability were assessed 42hrs post-transfection using the CytoTox 96 Nonradioactive Cytotoxicity Assay (Promega) or 3-(4,5-dimethyl-thiazol-2-yl)-2,5-diphenyl tetrazolium bromide (MTT) assay (Sigma), respectively, following the manufacturer’s protocol. Briefly, total LDH release was measured by lysing the cells with 1% Triton X-100 and absorbance was measured at 490 nm. MTT was added to cell culture media at a final concentration of 0.5 mg/mL and incubated for 45 minutes at 37°C. Following media removal cells were lysed with 100μl of Dimethyl sulfoxide (DMSO; Fisher Scientific) and absorbance was measured at 595 nm.

### RNA extraction and RT-qPCR

RNA was isolated from transiently transfected HEK293T or T98 cells using Trizol Reagent (Invitrogen). RNA was DNase treated using TURBO DNA-free Kit (Ambion), following the manufacturer’s instructions. cDNA was synthesized using random hexamer primers and the SuperScript III Reverse Transcriptase System (Invitrogen) following the manufacturer’s protocol. Quantification of construct transcript levels was performed using the 5FLAG (5’ GAT TAC AAG GAC GAC GAC GAC 3’) and 3HIS (5’ ATG GTG ATG GTG ATG ATG ACC 3‘) primers. Control reactions were performed using human β-actin forward (5′ TCG TGC GTG ACA TTA AGG AG 3′) and human β-actin reverse (5′ GAT CTT CAT TGT GCT GGG TG 3’) primers. qRT-PCR results were analyzed using the 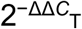 method (Livak & Schmittgen, 2001).

### Immunoblotting

HEK293T cells were washed with 1xPBS 48 hours post transfection and were lysed in 200μl radioimmunoprecipitation assay (RIPA) buffer (ThermoScientific) with 1X cOmplete Protease Inhibitors (Roche) for 15 min on ice. DNA was sheared by passage through a 21-gauge needle, lysates were centrifuged at 21,000×g for 15 min at 4°C, and the supernatant was collected. The protein lysate concentration was quantified using Pierce BCA Protein Assay Kit (ThermoScientific) and10μg of soluble protein lysates were separated on a 4-12% Bis-Tris gel (BioRad) and transferred to a nitrocellulose membrane. The remaining insoluble protein pellet was extracted in 2% SDS by incubating at 42°C for 3 hours with frequent repeated pipetting and incubated at room temperature overnight. Insoluble protein lysate was passed through a Dot Blot Apparatus (BioRad) onto a PVDF membrane. Membranes were blocked for 2 hours at room temperature in 5% dry milk in 1xPBS containing 0.05% Tween-20 (Sigma) and probed with anti-FLAG antibody (1:2,000), anti-myc antibody (1:1,000), 1C2 antibody (1:10,000), and anti-GAPDH antibody (1:5,000) overnight at 4°C in blocking solution. The membrane was incubated with species-specific HRP-conjugated secondary antibody (Amersham) in blocking solution, and bands were visualized with the ECL plus Western Blotting Detection System (Amersham). Quantification of protein expression was performed using Image J. For dot blot quantification, Myc antibody signal for empty vector transfections was used to perform background reduction. All protein levels are normalised to pure repeat expansion protein levels.

### Immunofluorescence (IF)

HEK293T cells were fixed 48 hours post-transfection with 4% paraformaldehyde (PFA; Sigma) in 1xPBS for 15min and permeabilized with 0.5% Triton X-100 (Sigma) in 1xPBS for 30 min. Cells were blocked in 1% Normal Goat Serum (NGS) for 30 minutes and incubated overnight at 4°C with 1C2 antibody (1:10,000) or anti-FLAG antibody (1:1,000), or for 1hr at 37°C with anti-myc antibody (1:1,000). Cells were incubated with AlexaFluor conjugated secondary antibodies for 1 hour at room temperature and were mounted with ProLong Gold Antifade (ThermoScientific). Representative images were taken using the ZEISS LSM 800 confocal microscope.

### UV melting

RNA oligonucleotides were purchased from IDT. Absorbance of each RNA substrate at 260nm was monitored between 25°C and 95°C, recorded at 1°C intervals. Three UV melting curves were generated per RNA substrate at a concentration of 2 μM in 1xDPBS without calcium or magnesium.

### Statistical Analysis

All statistical analyses were performed using GraphPad Prism 6.0 software. Statistical relationship of CCG•CGG interruptions and disease penetrance was calculated using Fisher’s exact test. Linear regression analyses were performed to assess the relationship between age of onset and repeat length or interruption number. All other statistical analyses were performed using unpaired two-tailed Student’s t-test or a one-way ANOVA with a Tukey’s multiple comparison test, as appropriate. Data are reported as mean ± SEM or mean ± SD.

## Acknowledgments

We thank study participants and their families for participating in our study. We thank Dr. M. Swanson, Dr. J.D. Cleary and members of the Ranum lab for helpful suggestions. This work was funded by R37 NS040389 (L.P.W.R.) and T32 training grant NS082168 (B.A.P.) and the National Ataxia Foundation.

## Author Contributions

B.A.P, H.K.S, Y.I., and L.P.W.R conceived the project. H.K.S., B.A.P., M.B.C., T.R., L.A.L., K.R, and Y.I. conducted experiments. H.K.S, B.A.P., Y.I., M.B.C, and K.R. analyzed data. S.H.S., C.G., T.A., J.W.D., L.J.S., L.F.H., J.E.N., and L.P.W.R. provided research participants. H.K.S., B.A.P. and L.P.W.R wrote the manuscript with input from all of the authors.

## Conflict of Interests

These authors disclose no competing interests.

---

**Table EV1.** Repeat length is not a reliable predictor of SCA8 disease status. Characteristics of participants are presented as mean ± SD, number and percentage of affected or asymptomatic individuals (%). To determine group effect, Fisher’s exact test was used for categorical variables and Mann-Whitney for non-parametric continuous variables. Average age of onset (Avg age onset) n=85. ns: not significant.

|  | Affected<br>n=111 | Asymptomatic<br>n=88 | p-value |
| --- | --- | --- | --- |
| Avg age onset (yrs) | 33.7 ± 19.7 | - |  |
| Females, n (%) | 57 (51.35) | 49 (55.68) | ns |
| Males, n (%) | 54 (48.65) | 39 (44.32) | - |
| Combined repeat # |  |  |  |
| Mean | 189 ± 219.9 | 188.5 ± 211.1 | - |
| Median | 113 | 98 | ns |
| Maximum | 1455 | 1000 | - |
| Minimum | 54 | 52 | - |

**Figure EV1.**
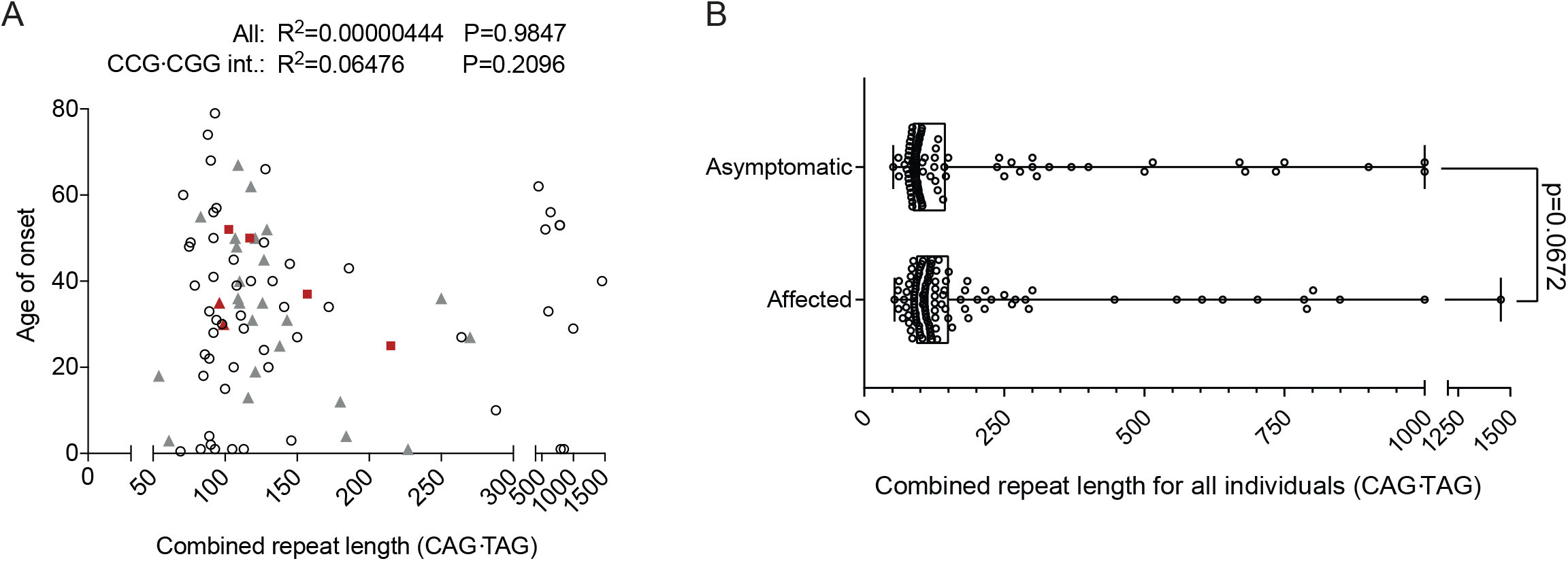
Repeat length is not a reliable predictor of age of onset or disease status in SCA8. A No correlation between length of combined repeat expansion and age of onset in SCA8 patients, n=85, p=0.9847 or in the subset of SCA8 patients with CCG•CGG interruptions, n=26, p=0.2096. Red squares indicate the average expansion size for individuals with two expanded alleles, individual allele repeat lengths: 137/177,110/320, 104/130, 96/109. Red triangles indicate the average expansion size for individuals with two expanded alleles and CCG•CGG interruptions: 84/114, 92/100. Grey triangles indicate individuals with CCG•CGG interruptions. B Allele length distribution of affected (n=111) and asymptomatic (n=88) expansion carriers, presented as minimum to maximum value, p=0.0672.

**Figure EV2.**
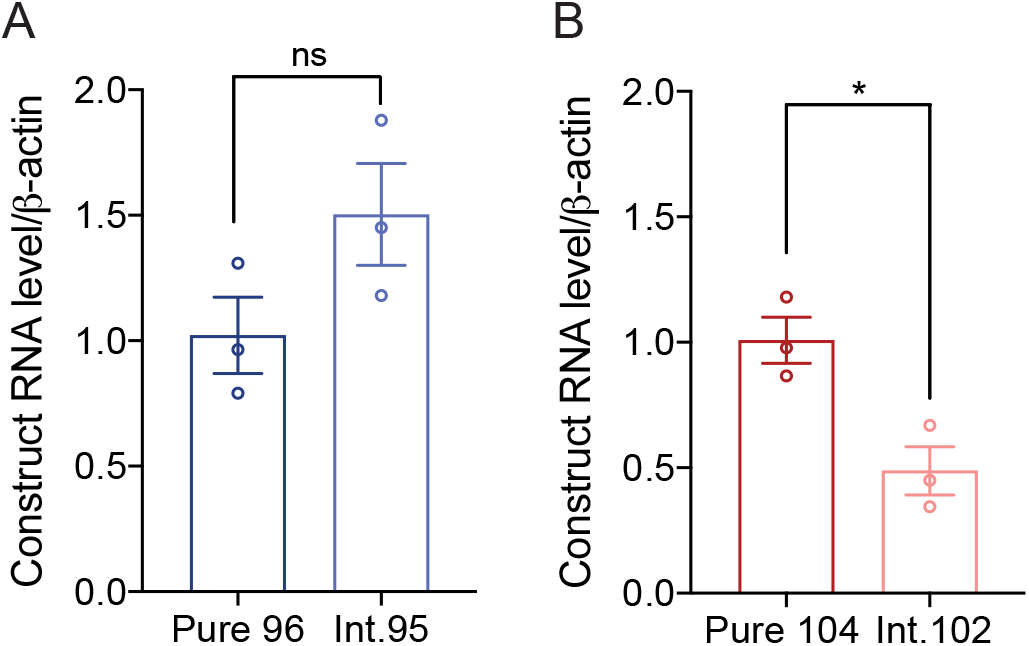
CGG interrupted SCA8 repeat tracts increase cellular toxicity independent of construct RNA levels. A qRT-PCR of Pure 96 and Int.95 construct transcript levels; n=3; p=0.1308, data presented as mean ± SEM. B qRT-PCR of Pure 104 and Int.102 construct transcript levels; n=3; p=0.0172, data presented as mean ± SEM.

**Figure EV3.**
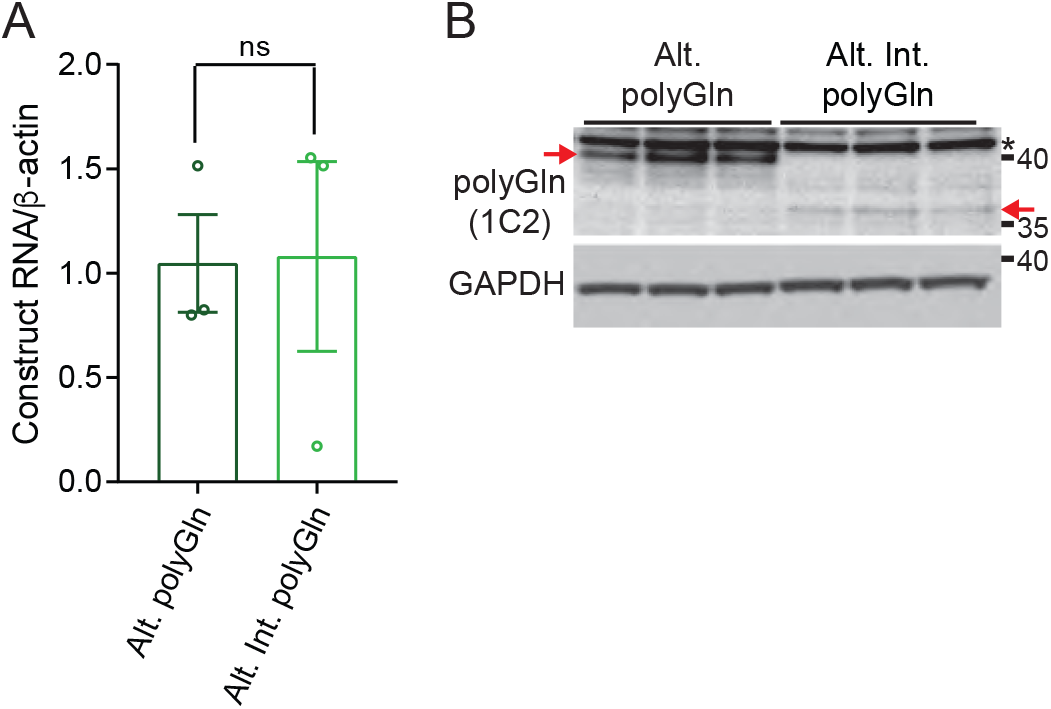
Arginine interruptions increase toxicity of polyGln proteins independent of RNA levels. A qRT-PCR of Alt. polyGln and Alt. Int. polyGln construct transcript levels, n=3; p=0.9516, ns: not significant, data are presented as mean ± SEM. B Western blot of polyGln proteins in HEK293T cells detected by 1C2 antibody from interrupted and pure CAA repeat tracts; red arrows indicate pure polyGln and polyGln(Arg) proteins. * The low levels of recombinant protein expressed for toxicity studies allows for polyGln containing TATA binding protein to be detected by 1C2 antibody giving a background band at ~40 kDa.

**Figure EV4.**
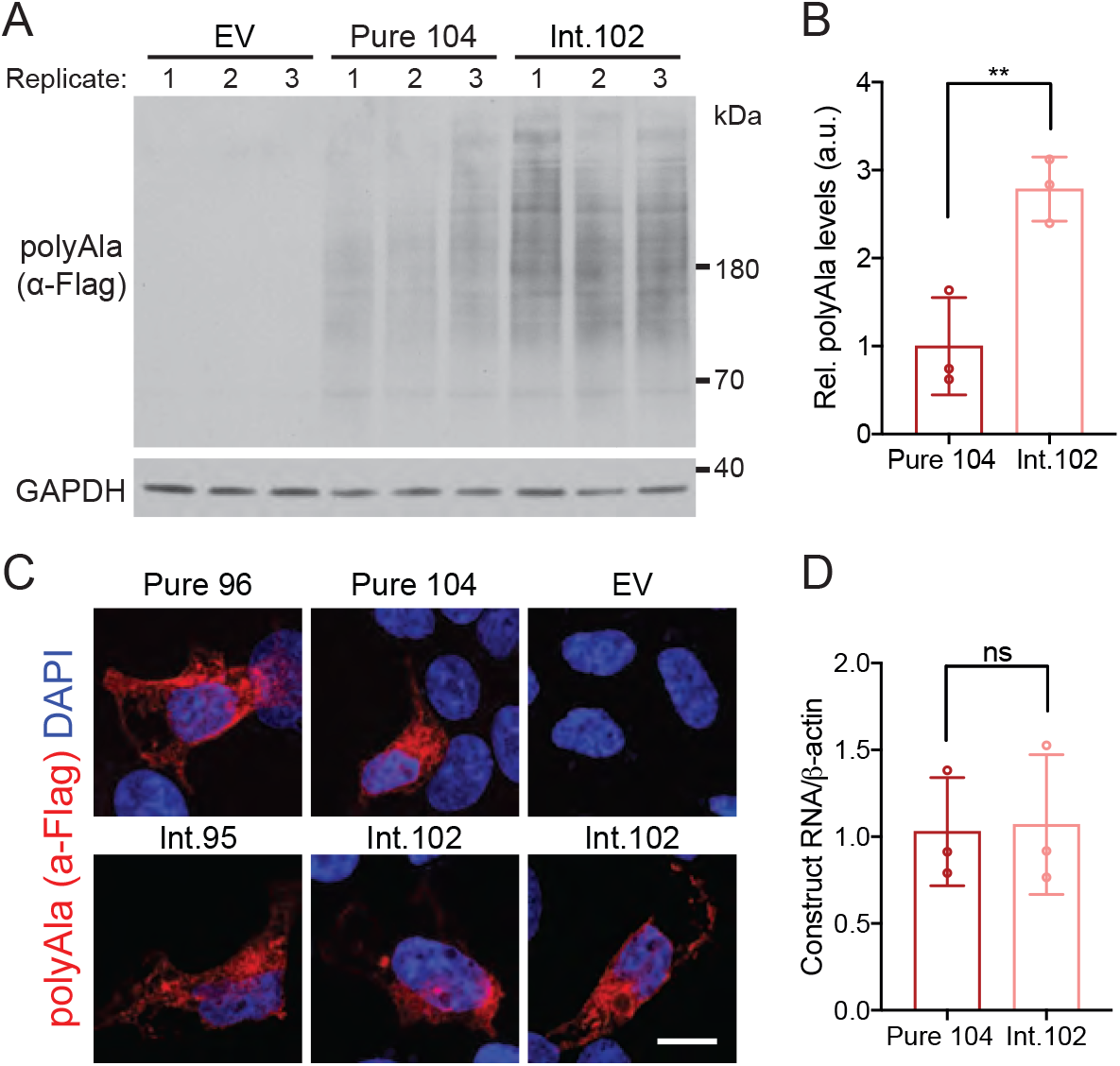
CGG interruptions increase levels of RAN polyAla expansion proteins. A, B Western blot (A) and densitometry quantification (B) of polyAla RAN proteins in HEK293T cells from interrupted (construct Int.102) and pure (construct Pure 104) CAG repeat tracts; EV: empty vector (pcDNA3.1-6S-3T), GAPDH as loading control; n=3, ** p<0.01, data presented as mean ± SD. C Immunofluorescence of RAN polyAla proteins in HEK293T cells; scale bar 10μm. D qRT-PCR of Pure 104 and Int.102 construct transcripts; n=3; p=0.8955, data presented as mean ± SEM.

**Figure EV5.**
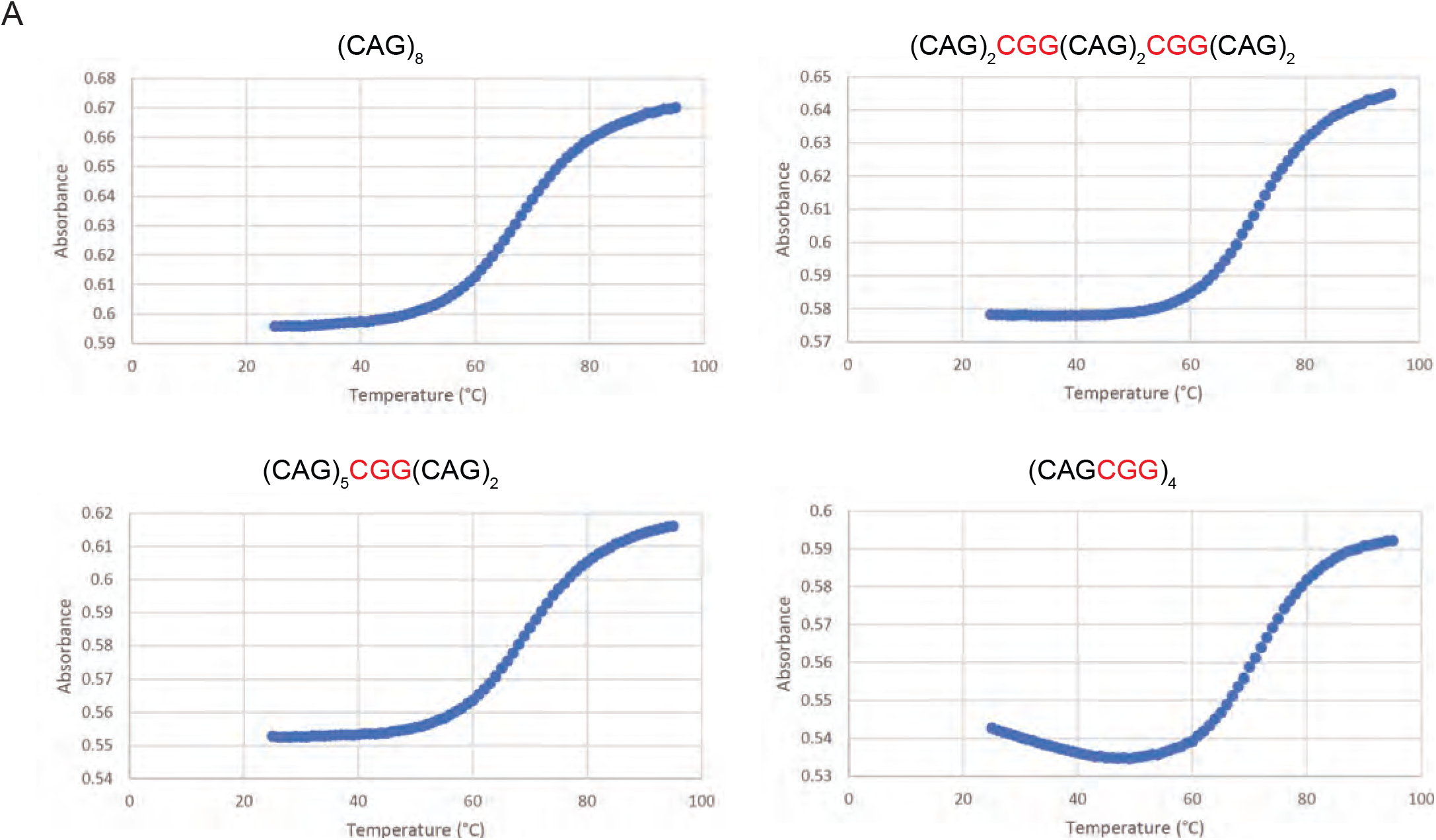
CGG interruptions increase stability of CAG repeat RNA hairpins. A Example UV melting absorbance curves (for Figure 5A) for pure and interrupted RNA oligos measured at 260nm monitored between 25°C and 95°C, recorded at 1°C intervals.

**Figure EV6.**
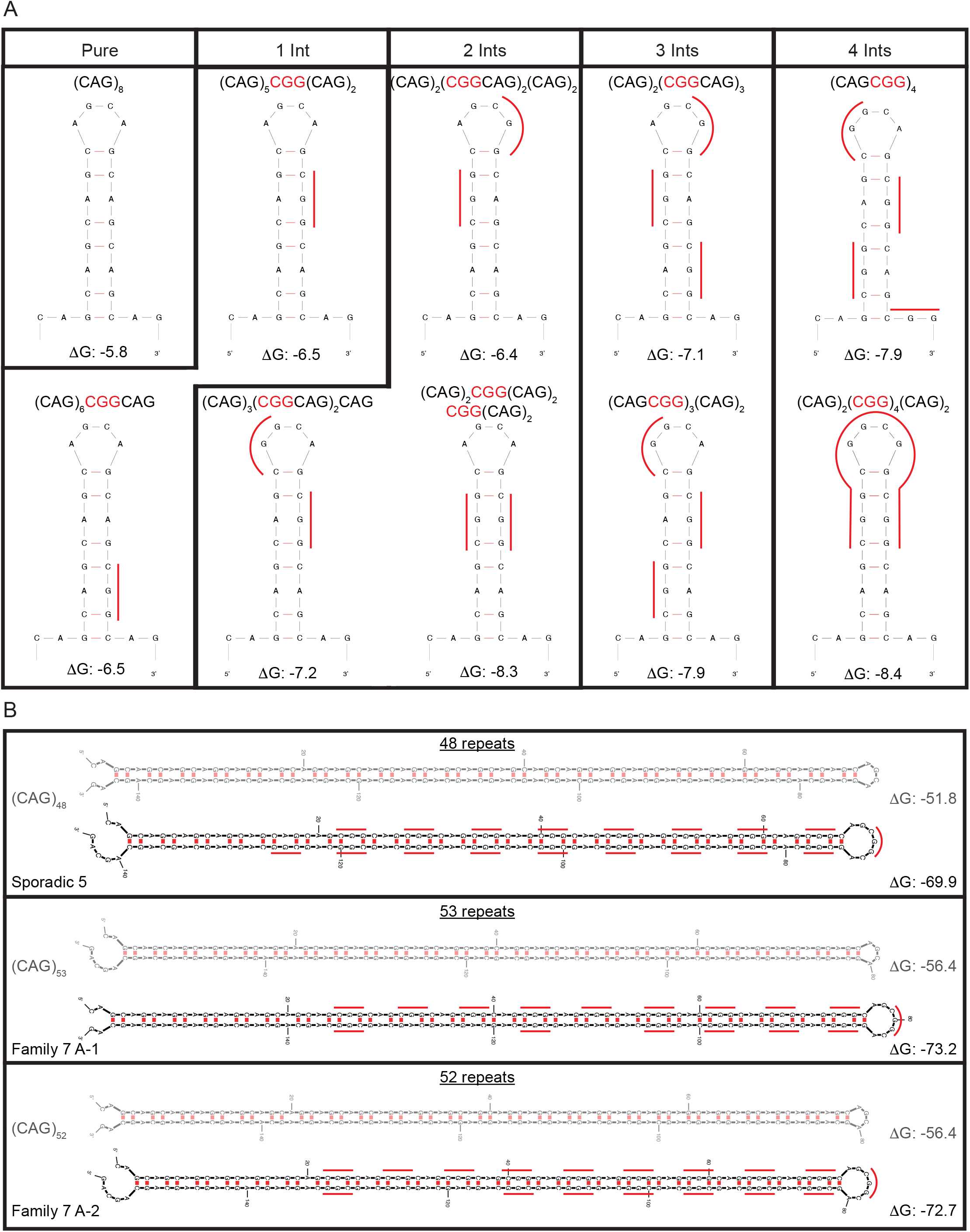
Predicted RNA structures for pure CAG repeat tracts and CGG interrupted CAG repeat tracts. A and B Predicted RNA hairpin structures from m-fold (Zuker, 2003) for pure and CGG interrupted CAG repeat tracts for Figure 5B (A) and Figure 5C (B). (B) Pure structures are shown in grey. For repeat tracts with multiple predicted hairpin structures, only the most stable structure is shown. Red lines alongside the structures indicate positions of CGG interruptions.

